# An amyloidogenic fragment of the SARS CoV-2 envelope protein promotes serum amyloid A misfolding and fibrillization

**DOI:** 10.1101/2024.04.25.591137

**Authors:** Asal Nady, Sean E. Reichheld, Simon Sharpe

**Affiliations:** Molecular Medicine Program, The Hospital for Sick Children, Toronto, Canada; Department of Biochemistry, University of Toronto, Toronto, Canada

## Abstract

SARS CoV-2 infection can affect a surprising number of organs in the body and cause symptoms such as abnormal blood coagulation, fibrinolytic disturbances, and neurodegeneration. Our study delves into the intricate pathogenic potential of a SARS-CoV-2 envelope protein peptide, shedding light on its implications for multi-organ effects and amyloid formation. Specifically, we focus on the peptide SK9 or ^54^SFYVYSRVK^62^ derived from the C-terminus of human SARS coronavirus 2 envelope protein. We demonstrate that SK9 containing peptides readily form classic amyloid structures consistent with predictions of amyloid aggregation algorithms. *In vivo*, overexpression of proteases such as neutrophil elastase during inflammation can potentially lead to C-terminal peptides containing SK9. We also demonstrate that SK9 can promote the fibrillization of SAA, a protein marker of acute inflammation. Our investigations reveal that the aromatic residues Phe2 and Tyr3 of SK9 play a pivotal role in its amyloidogenic function. We show that the primary sites of SK9-SAA binding lie in the amyloidogenic hotspots of SAA itself. Our results highlight two possible complications of SARS CoV-2 infection in individuals with hyper-inflammation either due to amyloids arising from SK9 containing peptides or SK9-induced AA amyloidosis.

## Introduction

Severe acute respiratory syndrome coronavirus 2 (SARS-CoV-2) is a highly transmissible and pathogenic coronavirus that emerged in late 2019^1,2^. The outbreak of this virus caused a global pandemic (Coronavirus disease 2019, COVID-19) with over 3.4 million deaths and 670 million infections (https://www.who.int/data/stories/the-true-death-toll-of-covid-19-estimating-global-excess-mortality). Although the majority of patients recover, a significant fraction experience post-acute sequelae (long-COVID), with neurological, respiratory and other symptoms persisting weeks to years post-infection^3,4^. Tissue damage in multiple organs, including the lung, brain, heart, and kidney, has been observed in patients with long-COVID, however little is known about the underlying molecular mechanisms^5–10^. Current estimates suggest about 10-40% of people infected with SARS-CoV-2 experience long COVID,^11,12^ with the majority of affected individuals also experiencing underlying chronic inflammation due to conditions such as diabetes, obesity, or cardiovascular disease with elevated levels of cytokines as IL-1β, IL-6, and TNF-α^13,14^. These cytokines are known to participate in the SARS-CoV-2 induced cytokine storm during acute infection, where they contribute to a vicious cycle of inflammation, and may also play roles in the onset and ongoing effects of long-COVID^13,15^.

Proinflammatory cytokines also promote the overexpression of an acute phase apolipoprotein called serum amyloid A (SAA)^16^. SAA is well characterized as a marker of inflammation and is expressed in the liver as part of the response to stress, infection, and injury^17–19^. During the acute phase of inflammation, SAA is primarily bound to circulating high-density lipoprotein (HDL) and has numerous proposed roles in innate immunity, pro-inflammatory response and tissue repair^20–23^. However, in individuals with chronic inflammation the levels of SAA are persistently high and there are increases in both the fraction of lipid-free SAA, and in non-hepatic expression of SAA^24–26^. In addition to exacerbating the underlying chronic inflammatory condition, long-term expression of SAA can result in the misfolding and aggregation of this protein, leading to systemic AA amyloidosis^27,28^As reported for other inflammatory conditions,^18,29–31^ serum levels of SAA have been positively associated with higher COVID-19 severity and mortality^19,32,33^ .

Due to its role in inflammatory processes, SAA has been proposed to play a role in COVID-19 associated hyperinflammatory syndrome^34^. Increased SAA levels during SARS-CoV-2 infection have also been shown to increase platelet adhesion, linking it with abnormal clotting and thrombosis in COVID-19 patients^35,36^. In patients with chronic inflammation, additional infection with COVID-19 can further increase their risk of AA amyloidosis due to covid-induced expression of proinflammatory cytokines and SAA, adding to their existing high levels^32,37^. The organs and tissues affected by systemic or localized AA amyloidosis are in many instances overlapping with those afflicted by COVID-19^38^. Therefore, it has been hypothesized that AA amyloidosis is an important factor causing systemic complications after COVID-19^39^.

In addition to acute phase proteins such as SAA, a heightened and prolonged inflammation can lead to increased expression of extracellular proteases such as neutrophil elastase (NE)^22,40,41^. A growing body of evidence correlates NE to the pathogenesis of COVID-19 disease^42^. Recent studies have shown that protease cleavage of several SARS-CoV-2 proteins (spike, ORF-10, ORF-6) results in the release of amyloidogenic peptides^43,44^. Adding to recent hypotheses linking viral infection and amyloid diseases,^38^ SARS-CoV-2 is able to promote amyloidogenesis of human proteins^45^. For example, the SARS-CoV-2 N protein accelerates fibrillization of human α-synuclein through an electrostatic interaction,^46^ and viral induction of surfactant protein amyloidosis in the lung has been proposed as a contributor to acute respiratory distress syndrome (ARDS) in COVID-19 patients^47^. Molecular dynamics simulations have been used to identify a putative interaction between SAA and a peptide from the C-terminus of the SARS-CoV-2 envelope (E) protein, ^54^SFYVYSRVK^62^ (SK9)^48^. SK9 binding was proposed to trigger SAA amyloid fibril formation by destabilizing the oligomeric state of SAA and increasing the population of aggregation prone monomeric SAA^48^. Interestingly, these amyloidogenic sequences may act as inflammatory stimuli that upregulate cytokines and the ACE2 receptor, increasing viral uptake by cells^44^.

In the current study we used Thioflavin T (ThT) fluorescence, circular dichroism (CD) spectroscopy and transmission electron microscopy (TEM) to show that the SK9 fragment of SARS-CoV-2 E forms amyloid fibrils. NMR spectroscopy and peptide arrays were used to show binding of SK9 to amyloidogenic regions located in helices 1 and 3 of SAA. SK9 binding was found to trigger formation of SAA amyloid fibrils *in vitro*, and key aromatic residues required for SK9 or SK9-induced SAA fibrillization were identified. These properties were largely retained by a longer 20-residue fragment of the E protein, suggests that proteolysis of E has the potential to release amyloidogenic peptides *in vivo*. Based on our findings, we suggest two possible mechanisms of amyloid formation that can lead to tissue damage following COVID-19 infection. Peptides derived from the E protein through proteolysis may self-assemble into amyloid fibrils, with potential cytotoxic effects. In addition, amyloidogenic E protein peptides may promote SAA aggregation, resulting in systemic AA amyloidosis. Both mechanisms depend on the increased inflammatory and immune response characteristic of COVID-19 to increase levels of NE and/or SAA at the sites of viral infection.

## Materials and Methods

### Peptides

The SARS-CoV-2 E protein derived peptides SFYVYSRVK (SK9), SAAVYSRVK (SK9AA), SFYVYSAVK (SK9RA), and LVKPSFYVYSRVKNLNSSRV (LV20) were obtained as HPLC-purified and lyophilized material from Genscript. All peptides were ordered with amidated C-termini and acetylated N-termini to mimic their position in the full-length E protein. For peptide array experiments the fluorophore-tagged peptide dansyl-GSGSGGSFYVYSRVK (dansyl-SK9GS) and a scrambled variant, dansyl-GSGSGGVKYSFRVYS (dansyl-SCR), were obtained from Genscript. The added GS residues provide a flexible linker to avoid fluorophore interference with SAA binding and fibril formation. Peptide solutions were prepared by weighing the lyophilized powder and dissolving it in the appropriate aqueous buffer at room temperature.

### Serum amyloid A expression and purification

Recombinant SAA2.1 (11.7 kDa) was expressed in BL21 (DE3) E. coli with an N-terminal His_6_ tag. SAA was purified from inclusion bodies using nickel-affinity chromatography run under denaturing conditions (6M GuHCl, 250mM NaCl, 50mM sodium phosphate pH 8.0) at room temperature. The purified SAA was dialyzed into a refolding buffer (400mM L-Arginine, 1.1M GuHCl, 21mM NaCl, 0.88mM KCl, 1mM EDTA, 55mM Tris pH 8.2) overnight at 4 °C followed by dialysis into a second stage refolding buffer (1M urea, 21mM NaCl, 20mM sodium phosphate pH 7.5) overnight at 4 °C. Refolded SAA was dialyzed into 20 mM Tris (pH 8.0) buffer containing 150mM NaCl, concentrated, and further purified using size exclusion chromatography (Superdex 200 10/300 GL). Under these conditions, SAA forms a stable, soluble octamer in the absence of phospholipids ^49^.

### Sequence analysis

The sequence of SARS-CoV-2 S, E, N, and M Proteins (Protein IDs: P0DTC2, P0DTC4, P0DTC9, P0DTC5) were subjected to the WALTZ algorithm (https://waltz.switchlab.org/) ^50^. The algorithms was set to high selectivity to avoid false positives. The sequence of the E protein (Protein ID: P0DTC4) was analysed using the AGGRESCAN server (http://bioinf.uab.es/aggrescan/) ^51^ to identify regions with high aggregation propensity. Expasy PeptideCutter (https://web.expasy.org/peptide_cutter/) ^52^ was used to predict neutrophil elastase cleavage sites within the E protein.

### Formation of lipid-bound SAA

Dimyristoyl phosphatidylcholine (DMPC) and dimyristoyl phosphatidylserine (DMPS) in chloroform were dried using nitrogen gas and residual solvent was evaporated by lyophilization. Lipids were suspended in water, subjected to several freeze-thaw cycles, lyophilized overnight, and then resuspended in 20 mM Tris buffer (pH 8.0) to form multilamellar vesicles (MLVs). MLVs were bath sonicated and extruded using 0.2 μm filtration membrane to form small unilamellar vesicles (SUVs), and stored at 37°C. The extruded SUVs were incubated with freshly purified SAA at mass ratio of 1:1 protein: lipid for 3 hours at 26°C to form lipid-bound SAA nanodiscs, as previously reported^49^. Nanodisc formation was confirmed using negative stain TEM. To test the effect of lipid headgroup charge, SAA nanodiscs were prepared with a varying ratio of DMPS: DMPC.

### Wide-angle X-ray scattering (WAXS)

Peptide solutions (4mg/ml in 20mM Tris, pH=8) were incubated for a week at 37 °C until mature fibrils formed. Amyloid Fibrils were lyophilized and tightly packed into a paste cell. An Anton Paar SAXSpace instrument with SAXSDrive software was used to collect scattering data. The sample to detector distance was 121.7579 mm. The data was collected at 25 °C. The one-dimensional data contain the scattered intensity as a function of the scattering vector magnitude *q* = (4π/λ)sinθ or *S* = (2π/λ)sinθ, where 2θ is the scattering angle and λ is the X-ray wavelength.

### Transmission electron microscopy

Samples were adsorbed to glow-discharged carbon-coated copper grids. Grids were subsequently washed with deionized water and stained with freshly prepared 2% uranyl acetate. Electron micrographs for all specimens were obtained with a Hitachi HT7800 transmission electron microscope operating at 80 keV. For single particle analysis of the negative stain TEM data, Relion (version 2)^53^ 2D classification was used for 200 particles, resulting in 3 class averages.

### Circular dichroism (CD) spectroscopy

Far-UV CD spectra were recorded using a Jasco J-810 spectropolarimeter. Samples were prepared contained 0.2 mg/mL protein in 20 mM Tris buffer, pH 8.0. Spectra were acquired at 4 °C in a 0.1 cm path length quartz cuvette and represent an average of three scans from 190-260 nm.

### Nuclear magnetic resonance spectroscopy

Solution ^1^H NMR data were obtained using a Bruker Avance III spectrometer with a ^1^H frequency of 600 MHz and a TXI triple resonance probe. NMR experiments were performed at 10 °C on samples containing 133 µM SAA in 20mM Tris buffer (pH 8.0). Peptide samples contained SK9, SK9AA or LV20 at 266 µM and mixed samples containing peptide and SAA were prepared at multiple peptide:protein ratios. NMRPipe^54^ was used to process the data and figures were generated using the Python libraries nmrglue and Matplotlib.

### Thioflavin T (ThT) fluorescence

A thioflavin T (Sigma-Aldrich) fluorescence assay was used to follow protein or peptide fibrilization using 96 well plates in a Synergy Neo2 multi-mode microplate reader. Aliquots of 10 µl of the incubated protein solution or peptide in powder form were added to 75 µL of PBS buffer pH 8.5 and 25 µL ThT pH 7.4. The plate was maintained at 37°C with 2 seconds of shaking before each read, with hourly measurements to 72 hours of total time. Excitation and emission wavelengths were 450 nm and 485 nm, respectively. The experimental conditions and controls (ThT and PBS buffer) were done in triplets.

### SAA peptide arrays

Peptide libraries were produced by automatic SPOT synthesis with standard Fmoc chemistry on continuous cellulose membrane supports using a Multipep 2 (CEM Corporation) peptide synthesizer. The presence of the peptides on the array was confirmed using Fast Green staining (0.1% Fast Green, 95% Ethanol) for 5 min. The stain was subsequently removed using 100% Methanol followed by PBS. The peptide binding assay was performed by rinsing the membrane with PBST (1x PBS, 0.1% TWEEN 20) three times for 5 mins. The membrane was then blocked for 1 hr with 5% milk powder in PBST at room temperature, then washed with PBST (2×5 mins) and PBS (5 mins). The array was incubated with 0.2 µM tagged peptides for 1 hr at 4 °C in dark. Unbound peptide was washed with PBST (3×5 mins). The interaction of supported SAA peptides with either dansyl-GSGSGGSFYVYSRVK or dansyl-GSGSGGVKYSFRVYS was determined (excitation at 335nm, emission monitored at 518nm) using an Odyssey imager (LI-COR Biosciences).

## Results

### Regions of the SARS-COV-2 E protein predicted to form amyloid fibrils

The WALTZ algorithm^50^ was used to identify putative amyloidogenic segments within the spike, envelope, nucleocapsid and matrix proteins from SARS-CoV-2 (Fig. 1, Supplementary Fig. S1). Given that amyloid formation by spike protein peptides has already been experimentally established ^44^, we focused on the E protein since it contains an exposed C-terminal domain (CTD) capable of interacting with serum proteins and potentially susceptible to proteolysis and release of peptides^56–58^. Supplementary Fig. S2 shows predicted sites of proteolytic cleavage of the E protein by NE, highlighting the likelihood of producing fragments containing the SK9 sequence previously shown to form amyloid fibrils *in silico*^48^. Only two segments with high amyloidogenic propensity were found within the exposed CTD: ^41^YCCNIVNVS^49^ and ^54^SFYVYSR^60^ (WALTZ). The 41-49 segment is unlikely to be released through proteolysis due to its proximity to the viral membrane, so we focused our study on peptides containing the ^54^SFYVYSR^60^ segment.

**Figure 1.**
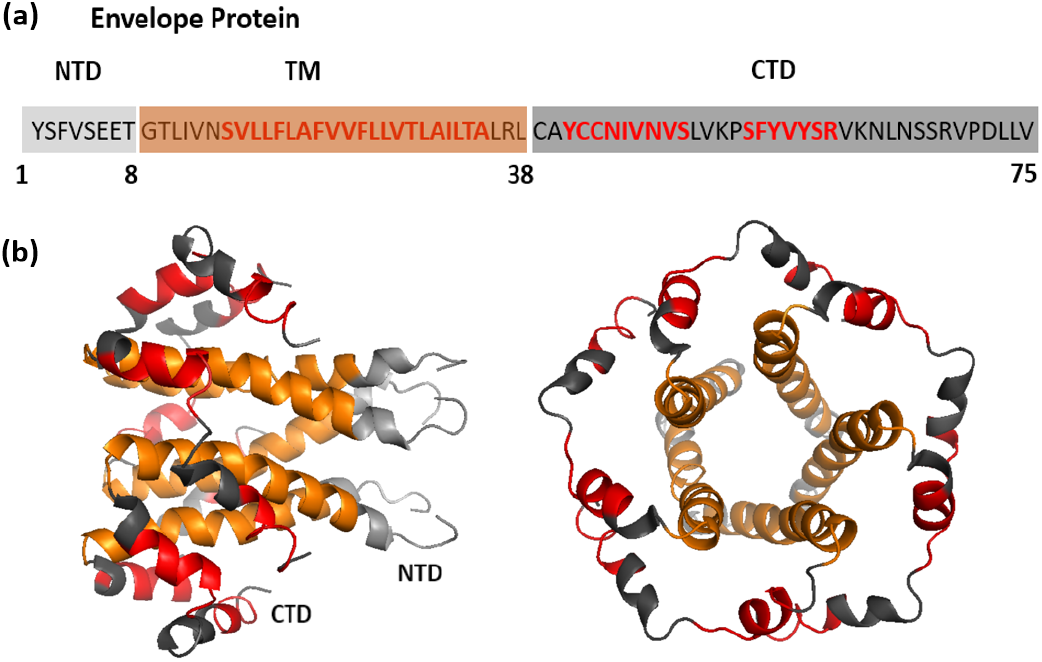
Predicted amyloidogenic regions of the SARS-CoV-2 envelope protein. (a) Amino acid sequence of SARS-Cov2 envelope protein. Sequences predicted to be amyloidogenic by the Waltz amyloid prediction algorithm (https://waltz.switchlab.org\)^50^ are highlighted in red. (b) Pentameric structure of the envelope protein (PDB: 5X29)^55^. The N-terminal domain is light grey, the C-terminal domain is dark-grey, and the transmembrane helices are orange.

### SK9 self-assembles into amyloid fibrils and two aromatic residues play a vital role

Upon incubation at relatively high concentrations (3.4 mM), TEM images show formation of amyloid-like aggregates of SK9 (Fig. 2). At shorter incubation times and lower peptide concentrations (0.4 mM), fibrillar assemblies composed of paired twisted filaments were observed, with small spherical objects decorating the surface of the fibrils (Fig. 2a). Class averaging of these beaded structures revealed a hollow centre with lower electron density within the bead-like structures suggesting the formation of protofibrillar oligomers in combination with mature fibrils under this condition. Similar annular protofibrils have been observed for numerous other amyloid peptides and are frequently associated with cytotoxicity ^59–61^.

**Figure 2.**
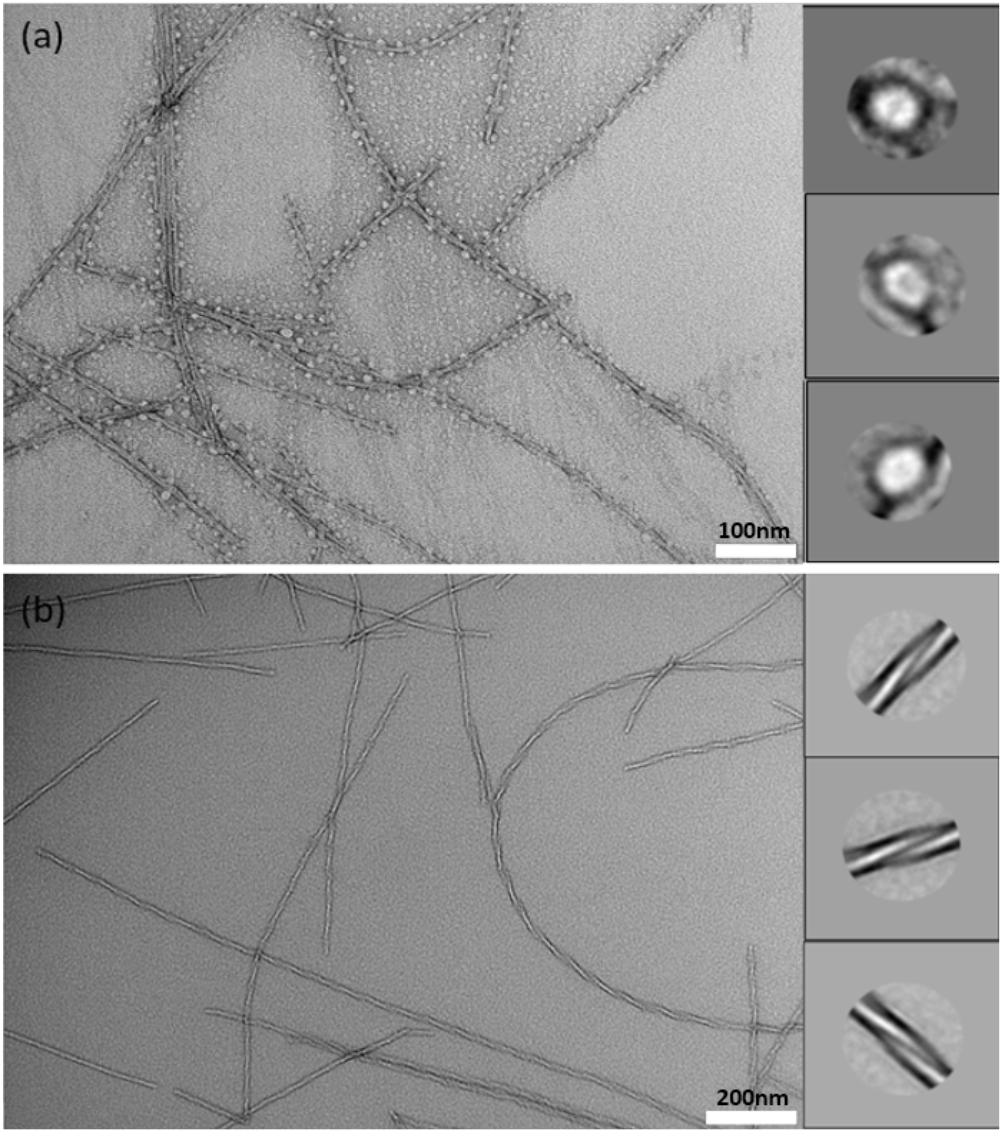
SK9 forms twisted multi-filament fibrils and protofibrillar oligomers. Negative stain TEM images of (a) SK9 fibrils formed by incubating 0.4 mM peptide in 20 mM pH 8 Tris buffer at 37 °C for 3 days (magnification=70000x), and (b) SK9 fibrils formed after 1 month of incubation at 4 °C. 3.4 mM peptide was incubated in 20 mM pH 8 Tris buffer (magnification = 30000x). Class averages shown on the right were obtained through averaging of 200 particles in each case, showing detail of the protofibrillar oligomers (a) and mature fibrils (b).

Longer incubation and higher peptide concentrations resulted in samples containing only mature amyloid fibrils, suggesting that either the protofibrillar aggregates are metastable and convert to mature fibrils over time, or that the fibrillar form is kinetically favored at high peptide concentrations. Class averaging of the mature SK9 fibrils (Fig. 2b) further reveals internal details of the twisted paired-filament structure, which is consistent with an amyloid-like internal structure. The low degree of ThT binding to SK9 fibrils is similar to prior reports of fibrils formed by the S protein ^44^. The amyloid nature of SK9 was also confirmed using WAXS (Supplementary Fig. S3) where SK9 fibrils give rise to a Bragg peak at 4.7 Å which corresponds to the interstrand hydrogen bond distance typically observed in amyloid fibril structures^62^.

To test the role of specific amino acids in SK9 fibril formation, we replaced the first two aromatic amino acids with alanine (SK9AA; SAAVYSRVK), a sequence that is not predicted by the WALTZ algorithm to form amyloid^50^. In a second modified peptide an arginine to alanine substitution was made to reduce the ability to form stabilizing intermolecular cation-π interactions (SK9RA; SFYVYSAVK)^63,64^. A 20 amino acid long sequence containing additional naturally occurring residues flanking SK9 was used to test the amyloid propensity of longer segments of the E protein containing an amyloidogenic core sequence (LV20; LVKPSFYVYSRVKNLNSSRV).

The modified SK9 peptides were incubated in pH 8 buffer, 37 C, for two weeks and imaged by negative stain TEM (Fig. 3). Substitution of aromatic residues in SK9 (Phe2 and Tyr3) completely abolished fibrillization, with only some non-specific aggregation observed. Similarly, SK9RA also showed substantially reduced fibril formation relative to the wild-type sequence, with a small number of short, irregular fibrils observed. The longer 20-residue peptide LV20 retained the ability to form fibrils, although there were notable differences from SK9. LV20 fibrils displayed a different morphology and appeared as needle-like crystalline assemblies accompanied by protofibrillar oligomers (Fig. 3d). The peptides were monitored for *in vitro* amyloid formation using ThT and SK9 resulted in the highest signal (Supplementary Fig. S4). Further, CD spectroscopy on SK9 and variant peptides (Supplementary Fig. S5) shows that all peptides are disordered in solution prior to aggregation, with the highest degree of disorder is seen for the non-fibrillogenic SK9AA.

**Figure 3.**
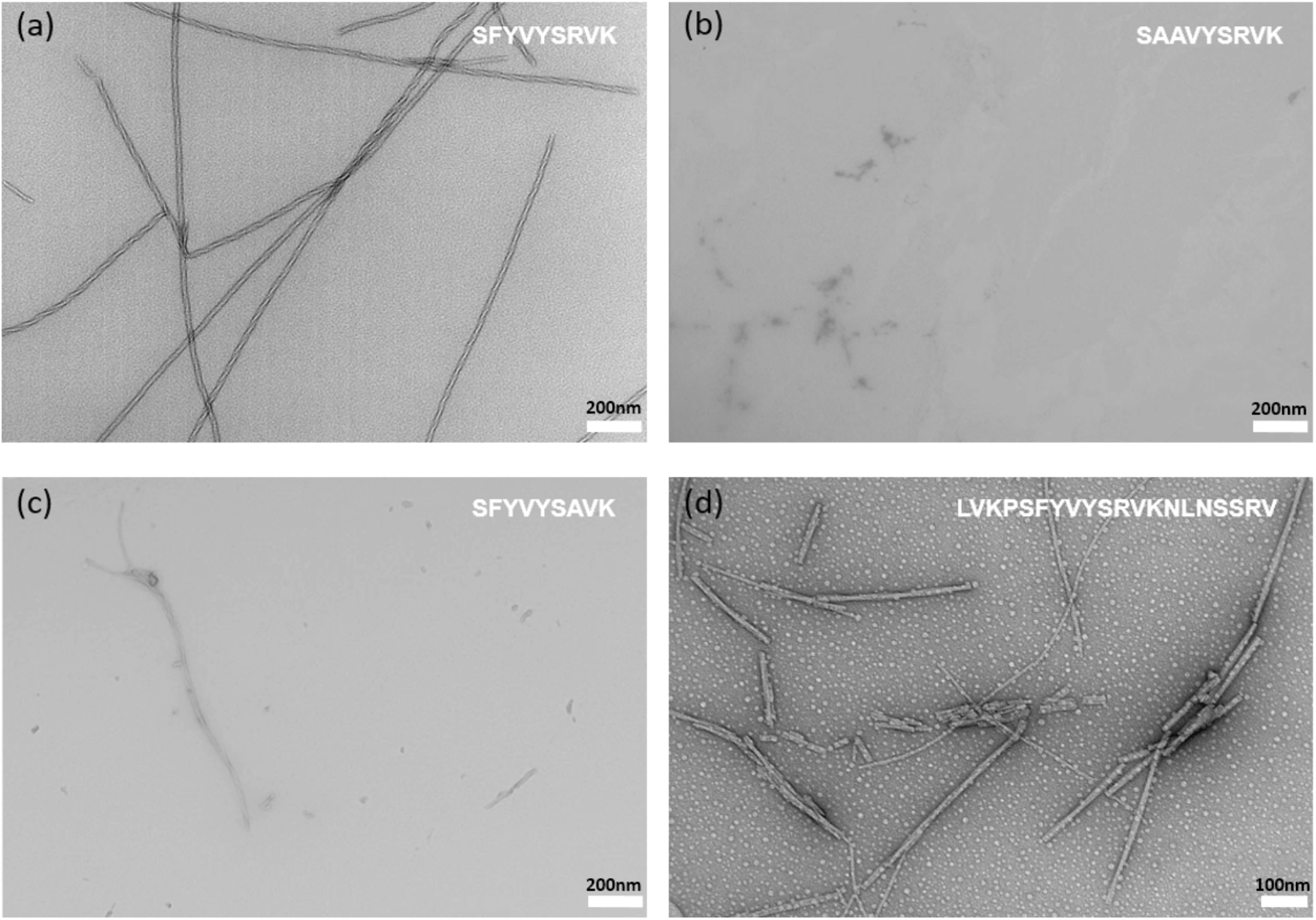
Amyloid fibril formation by SK9 peptide variants. Negative stain TEM images of (a) wild-type SK9 fibrils (magnification = 30000x), (b) SK9AA (magnification = 30000x), (c) SK9RA (magnification = 30000x) and (d) LV20 (magnification = 50000x). All peptides were incubated at 37 °C for two weeks prior to imaging.

### SK9 promotes the fibrilization of lipid-free SAA

To test the hypothesis that SK9 might trigger amyloid fibril formation by human SAA ^48^, we incubated SK9 with recombinant SAA octamers or SAA-lipid nanodiscs for 2 days at 37 °C. The latter were used to mimic the predominant HDL-bound form of SAA found during acute phase inflammation. ThT fluorescence and TEM microscopy were used to track the formation of amyloid fibrils by SAA at varied peptide: protein ratios (Fig. 4). A very low ThT fluorescence intensity was observed after addition of SK9 to SAA-lipid nanodiscs, regardless of DMPC:DMPS ratio, while addition of SK9 to lipid-free SAA gave a high ThT signal following 12 hrs of incubation. Under these conditions, SAA alone did not form amyloid fibrils, and any ThT signals expected for SK9 (Fig. S4) would be much weaker than observed, indicating that the signal arises from SAA fibrillization in the presence of peptide. ThT data obtained at varied ratios of SAA:SK9 (Fig. 4c), shows a peptide-dependent increase in fibrillization, further supporting a direct role for this E protein fragment in initiating formation of SAA amyloid fibrils. TEM data (Fig. 4a) show that SK9-induced SAA fibrils have a relatively irregular morphology that is distinct from the twisted pair fibrils formed by SK9 alone.

**Figure 4.**
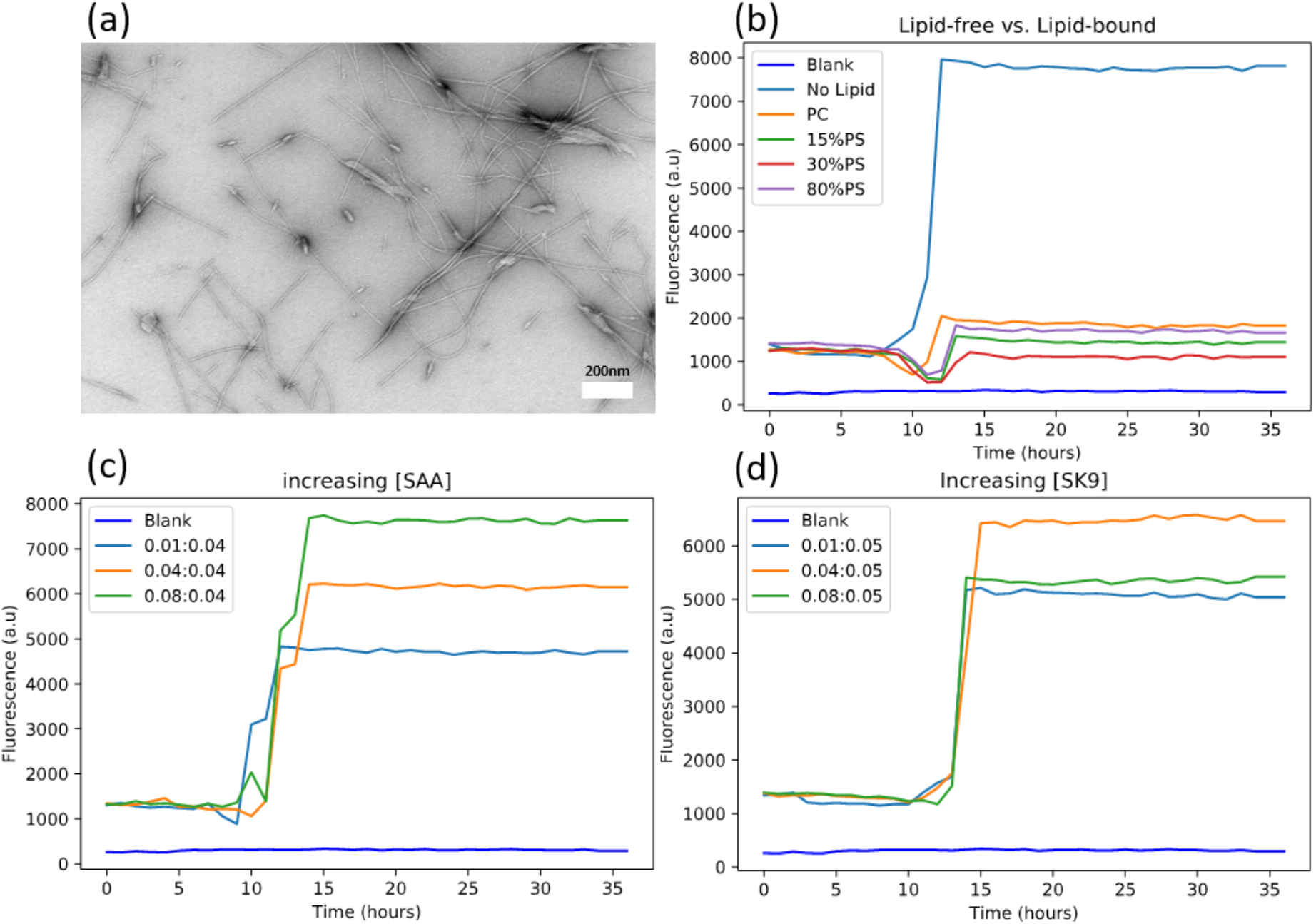
SK9 amyloid fibril formation by lipid-free SAA. (a) Negative-stain TEM image of fibrils formed after incubation of lipid-free SAA with SK9 peptide (magnification=30000x). (b) ThT fluorescence emission at 485nm recorded for 36 hours after addition of SK9 to samples containing lipid-free SAA, SAA-lipid nanodiscs made with either 100%DMPC, 15%DMPS:85%DMPC, 30%DMPS:70%DMPC, or 80%DMPS:20%DMPC. (c) ThT data following addition of increasing amounts of SAA to a sample containing 40 ug/mL SK9. (d) Shows similar data for samples containing 50 ug/mL SAA following addition of increasing amounts of SK9. All ThT fluorescence data were recorded at 37 °C.

### SK9 binds to lipid-free SAA prior to inducing fibril formation

To determine whether the SK9 peptide binds to SAA prior to initiating fibrillization, we performed trypsin digestion of SAA following incubation with SK9 or derivative peptides (Fig. 5). At incubation times shorter than the lag phase for fibril formation (data for 30 minutes are shown), addition of SK9 reduced SAA cleavage by trypsin (Fig. 5a). After 1 day of incubation amyloid fibrils were formed, resulting in significant protection from cleavage due to the stability of the fibril structure (Fig. 5b). SK9RA and LV20 also protected SAA from trypsin after 30 min incubation, although complete proteolysis was observed after 30 min digestion, suggesting a weaker interaction than observed for SK9. SK9AA (Fig. 5d) did not prevent trypsin digestion of SAA, consistent with the fibrillization results in Fig. 4. Bovine serum albumin (BSA) was used as a control, with SK9 incubation having no effect on tryptic digestion.

**Figure 5.**
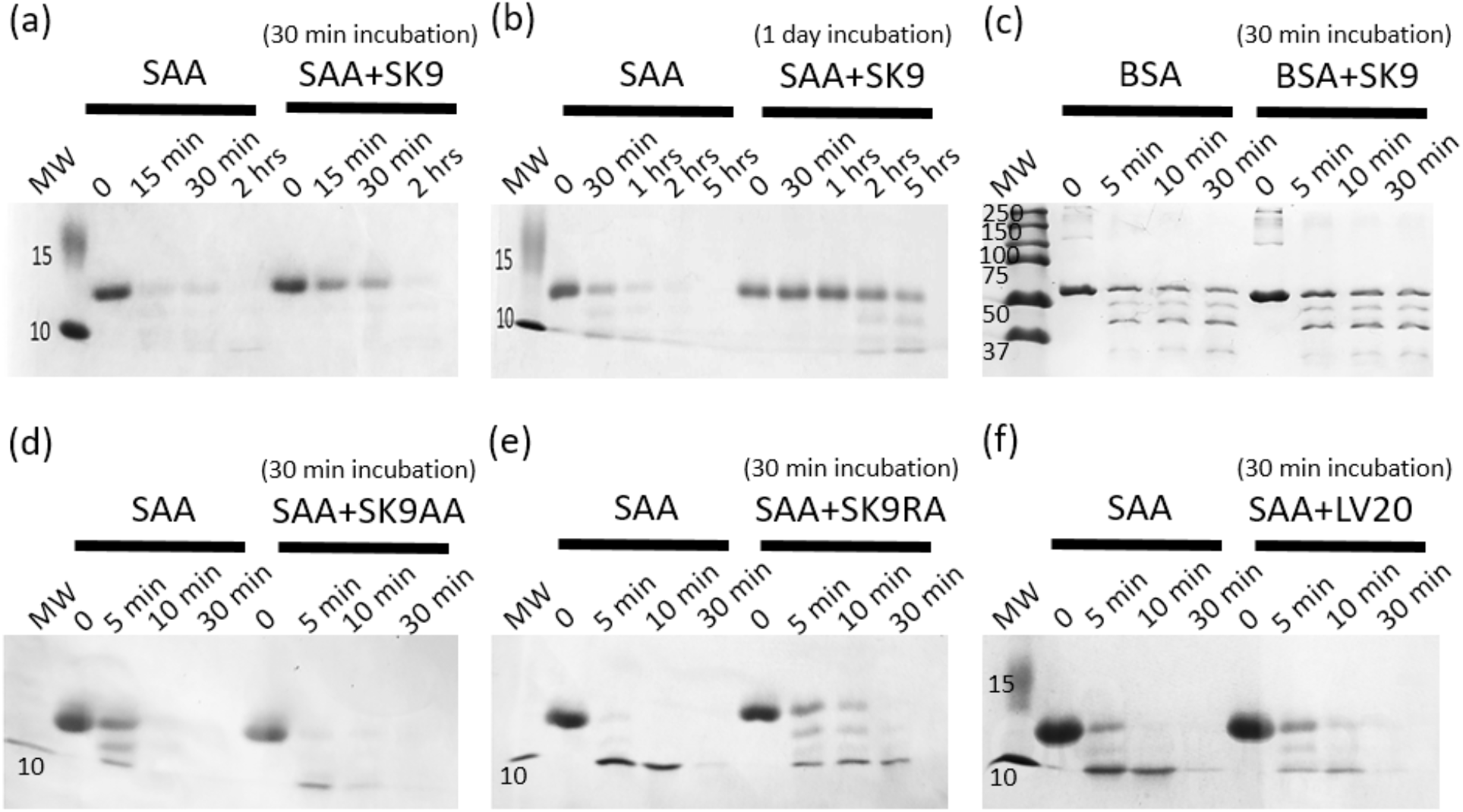
SK9 protects lipid-free SAA from trypsin proteolysis. SDS-PAGE analysis of SAA alone and in the presence of wild-type SK9 peptide following (a) 30 minutes of protein-peptide incubation and (b) 1 day of incubation at 37 °C. (c) Trypsin digestion of BSA and BSA incubated with SK9 for 30 minutes prior to digestion treatment. Trypsin digestion SDS-PAGE of SAA alone and in the presence of (d) SK9AA, (e) SK9RA, or (f) LV20. The trypsin reaction time is shown above each lane.

SAA protein and peptides were only incubated for a short period of 30 minutes at 37 °C to assay for protection due to binding and not fibrillization. Figures 5a,d-f show that the addition of wild-type SK9 results in the highest level of protection against trypsin digestion relative to the other peptide variants, followed by SK9RA or LV20. Interestingly, no protection against trypsin digestion is observed in SK9AA. Further, prolonged incubation of lipid-free SAA with SK9 (Fig. 5b) results in even greater protection likely due to the formation of amyloid at the 1 day time point which exceeds the 12 hr lag phase. We also performed the assay using bovine serum albumin (BSA) to determine if the protective effect due to the presence of the SK9 peptide is specific to SAA. Figure 5c shows that the presence of SK9 has no effect on the digestion of BSA by trypsin. These data were further confirmed using ^1^H NMR to monitor free vs bound SK9 peptides in the presence of increasing concentrations of SAA (Fig. S6), with only SK9AA retaining sharp methyl signals associated with free peptide.

To identify the specific segments of SAA that interact with SK9, a peptide microarray was synthesized containing 49 overlapping 10-residue peptides covering the entire SAA sequence. The array was probed with either dansyl-SK9GS or a scrambled control peptide (dansyl-SCR). In each case, the dye was attached using an extended GS sequence as a flexible linker. Figure 6a shows the presence of three primary hotspots for SK9 binding, while very little signal is observed in the control experiment (Fig. 6b), suggesting specificity of the native SK9 sequence for interaction with SAA. The strongest binding region is located in SAA helix 1 and contains a well-established amyloidogenic hexapeptide, SFFSFL^28^. A second interaction hotspot in helix 3 contains the sequence WAAEVI, which is known to form amyloid fibrils, albeit less readily than helix 1 peptides^28,65^. This is consistent with the greater degree of SK9 binding to helix 1. Further, both of these SAA segments are predicted to be aggregation prone (Supplementary Fig. S7). An additional interaction hotspot in helix 4 lacks amyloid or aggregation-prone sequences and is therefore less likely to play a role in fibrillization.

**Figure 6.**
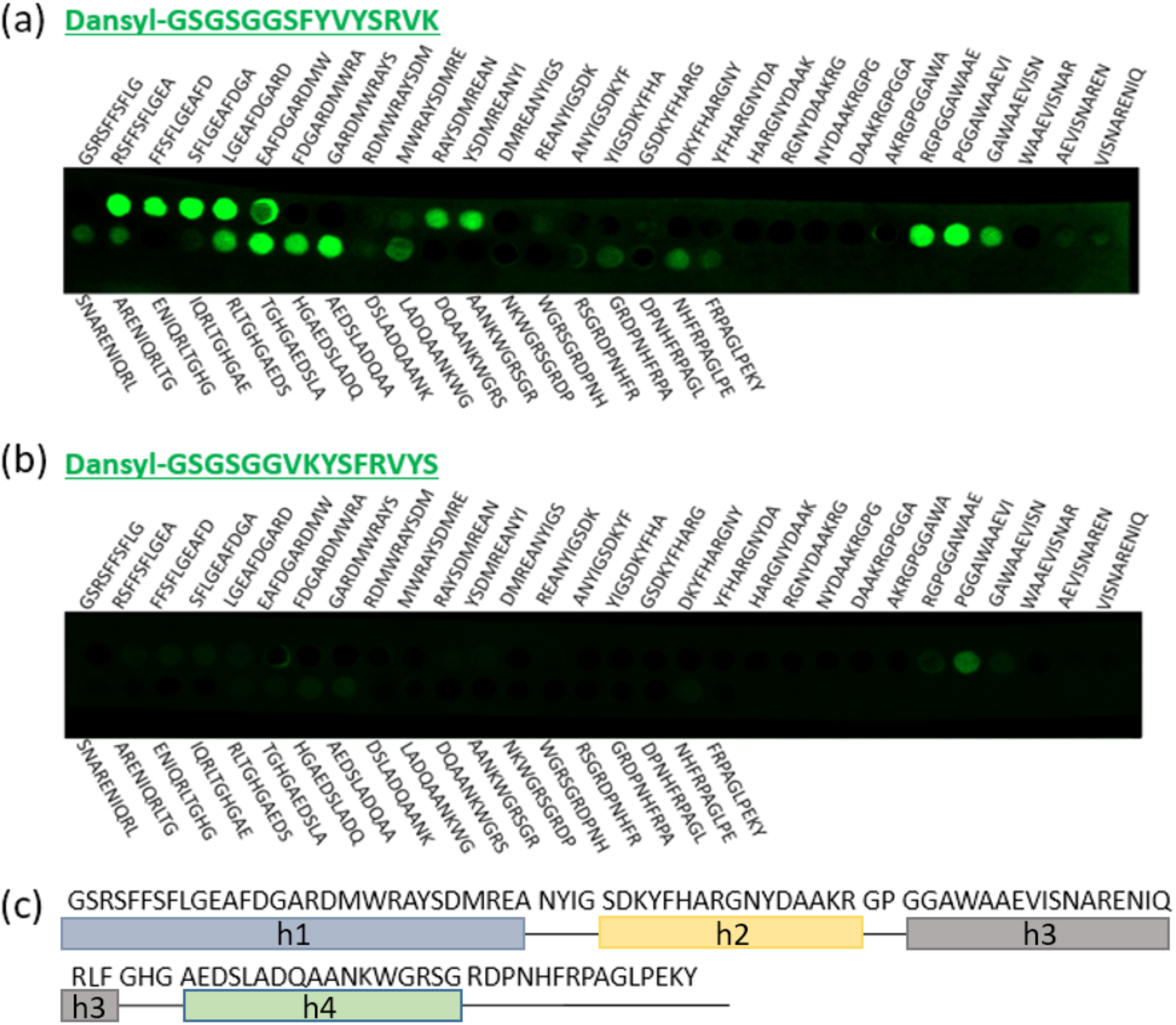
SK9 binds the amyloidogenic sequences of SAA. A peptide microarray was created using overlapping 10-residue segments from the full-length human SAA2 sequence, with the sequences as indicated. Each peptide has an 8 residue overlap with the adjacent peptides. Fluorescence images were obtained after incubating the microarray membrane with (a) dansyl-SK9GS or (b) the negative control peptide dansyl-SCR. (c) Boxed regions show the location of α-helices along the primary sequence of SAA.

## Discussion

SARS-CoV-2 is predominantly a respiratory disease affecting the lungs yet can have wide-ranging impacts throughout the body such as abnormal blood coagulation, fibrinolytic disturbances, neurodegeneration in the brain, and AA amyloidosis ^66–69^. While some recent studies have proposed putative links between SARS-COV-2 proteins and amyloidosis ^43,44^, including induction of amyloid formation by human proteins such as surfactant protein-C ^47^, *α*-synuclein ^46^, and SAA ^48^, relatively little is known about the underlying molecular mechanisms that lead to such complications. Similar links between viral infection and amyloidosis have been made for other human viruses, including HIV, HSV-1 and Influenza A, both as a direct consequence of aggregation-prone viral proteins and as a secondary effect of inflammation ^38^.

Similar to previous reports of amyloidogenic peptides derived from the SARS-CoV-2 S protein and ORF proteins ^43,44^, we have demonstrated *in vitro* amyloid fibril formation by a sequence within the viral E protein. This SK9 fragment was initially identified in a prior computational and molecular dynamics study and proposed to interact with SAA as a potential trigger of AA amyloidosis ^48^. Our data show that the SK9 sequence is able to rapidly assemble into amyloid fibrils that require aromatic residues for stability, likely due to the formation of π-π and cation-π interactions similar to those seen for other peptide fibrils^70,71^. Extending the length of the peptide did not impair fibrillization, suggesting that longer E protein fragments would also be amyloidogenic. Annular protofibrillar assemblies were also formed by SK9 and the longer SK20 peptide. Such species have been associated with the cytotoxicity of other amyloid proteins,^59,61^ making this a potential mechanism for increased tissue damage and inflammation in COVID-19.

We have shown that SK9 and LV20 are capable of forming long-lived interactions with helices 1 and 3 of SAA, initially protecting the protein from tryptic digestion and subsequently resulting in formation of SAA amyloid fibrils. This interaction is similar to the one predicted by an earlier MD study ^48^, and required the same SK9 sequence elements as formation of peptide-only amyloid fibrils. *In vivo*, the amyloidogenic sequences in SAA are likely protected from misfolding through native packing which includes either protein-protein or protein-lipid interactions. In the absence of lipids, SAA self-associates and forms either hexameric or octameric oligomers to protect its amphipathic helices and prevent aggregation ^28,72^. For SAA to form fibrils, it must first dissociate from its multimeric alpha helical form and form extended β-sheet structures^73^. Our data suggest that interaction of SK9 with the amyloidogenic hotspots of SAA is sufficient for oligomer destabilization and subsequent aggregation.

Notably, SK9 did not promote fibrillization of SAA bound to lipid nanodiscs that mimic the naturally occurring HDL-bound form of the protein. The most likely SK9 binding site in helix 1 of SAA is known to be a key HDL-binding site, which could explain why lipid-bound SAA is largely protected from SK9 interaction and amyloid formation^74^. This observation suggests that the presence of phospholipids protects the hydrophobic surfaces of SAA that may play a role in its fibrillization and that conditions which increase the levels of lipid-free SAA *in vivo* such as chronic inflammation may increase the risk of amyloidosis. SAA is normally bound to HDL in the serum during the acute inflammatory response, interacting with the lipoprotein surface via amphipathic helices ^23,28,75^. However, the levels of lipid-free SAA in serum have been shown to increase *in vivo* during periods of prolonged inflammation, as might be expected in long-term viral infection or long-COVID^26^. This is also consistent with the higher risk of secondary complications from COVID-19 in patients with underlying medical conditions that result in a long-term elevation of SAA in multiple tissues.

The ability of SK9 to interact with multiple segments of SAA which do not share sequence similarity but instead share a predicted potential for aggregation or formation of extended beta-strand structures suggests that this peptide may be capable of binding to many other amyloid forming sequences. In combination with the reported amyloidogenic potential of several regions of the SARS-CoV-2 S protein and viral accessory proteins ^43,44^, there is a strong potential for amyloidosis to contribute to the systemic damage caused by COVID-19.

In conclusion, our results suggest two possible pathways of fibrillization due to SARS-CoV-2 infection that are both fueled by a heightened innate inflammatory response. 1) Overexpression of extracellular proteases in response to inflammation leading to the production of amyloidogenic peptides from the CTD of the viral E protein. 2) Overexpression of SAA during the inflammatory response, in particular the increased levels of lipid-free SAA that occur in chronic inflammation, with subsequent interactions with the E protein promoting AA amyloidosis. Each of these pathways would result in increased local inflammation, potentially contributing to the long-term effects of COVID-19 infection.

## Acknowledgements

This work was funded by a Discovery Grant from the Natural Sciences and Engineering Research Council of Canada (held by S.S.), and a SickKids Restracomp scholarship (to A.N.). This work made extensive use of infrastructure provided by the Structural Biophysics Core facility and the Nanoscale Biomedical Imaging Facility at the SickKids Research Institute.

## REFERENCES

1. Lai, C. C., Shih, T. P., Ko, W. C., Tang, H. J. & Hsueh, P. R. Severe acute respiratory syndrome coronavirus 2 (SARS-CoV-2) and coronavirus disease-2019 (COVID-19): The epidemic and the challenges. International Journal of Antimicrobial Agents vol. 55 105924 Preprint at 10.1016/j.ijantimicag.2020.105924 (2020).

2. Wang, C., Horby, P. W., Hayden, F. G. & Gao, G. F. A novel coronavirus outbreak of global health concern. The Lancet vol. 395 470–473 Preprint at 10.1016/S0140-6736(20)30185-9 (2020).

3. Raveendran, A. V, Jayadevan, R. & Sashidharan, S. Diabetes & Metabolic Syndrome : Clinical Research & Reviews Long COVID : An overview. Diabetes & Metabolic Syndrome: Clinical Research & Reviews 15, 869–875 (2021).

4. Wu, Z. & McGoogan, J. M. Characteristics of and Important Lessons From the Coronavirus Disease 2019 (COVID-19) Outbreak in China. JAMA 323, 1239–1242 (2020).

5. Kommoss, F. K. F. et al. The Pathology of Severe COVID-19-Related Lung Damage. Dtsch Arztebl Int 500–506 (2020) doi:10.3238/arztebl.2020.0500.

6. Feng, G. et al. Covid-19 and liver dysfunction: Current insights and emergent therapeutic strategies. Journal of Clinical and Translational Hepatology vol. 8 18–24 Preprint at 10.14218/JCTH.2020.00018 (2020).

7. Douaud, G. et al. SARS-CoV-2 is associated with changes in brain structure in UK Biobank. Nature 604, 697–707 (2022).

8. Meinhardt, J. et al. Olfactory transmucosal SARS-CoV-2 invasion as a port of central nervous system entry in individuals with COVID-19. Nat Neurosci 24, 168–175 (2021).

9. Topol, E. J. COVID-19 can affect the heart. Science vol. 370 408–409 Preprint at 10.1126/science.abe2813 (2020).

10. Głowacka, M., Lipka, S., Młynarska, E., Franczyk, B. & Rysz, J. Acute kidney injury in COVID-19. International Journal of Molecular Sciences vol. 22 8081 Preprint at 10.3390/ijms22158081 (2021).

11. Taquet, M., Dercon, Q. & Harrison, P. J. Six-month sequelae of post-vaccination SARS-CoV-2 infection: A retrospective cohort study of 10,024 breakthrough infections. Brain Behav Immun 103, 154–162 (2022).

12. Chen, C. et al. Global Prevalence of Post-Coronavirus Disease 2019 (COVID-19) Condition or Long COVID: A Meta-Analysis and Systematic Review. Journal of Infectious Diseases 226, 1593–1607 (2022).

13. Low, R. N., Low, R. J. & Akrami, A. A review of cytokine-based pathophysiology of Long COVID symptoms. Front Med (Lausanne) 10, 1011936 (2023).

14. Buicu, A. L., Cernea, S., Benedek, I., Buicu, C. F. & Benedek, T. Systemic inflammation and COVID-19 mortality in patients with major noncommunicable diseases: Chronic coronary syndromes, diabetes and obesity. Journal of Clinical Medicine vol. 10 1545 Preprint at 10.3390/jcm10081545 (2021).

15. Hasanvand, A. COVID-19 and the role of cytokines in this disease. Inflammopharmacology vol. 30 789–798 Preprint at 10.1007/s10787-022-00992-2 (2022).

16. De Buck, M. et al. The cytokine-serum amyloid A-chemokine network. Cytokine and Growth Factor Reviews vol. 30 55–69 Preprint at 10.1016/j.cytogfr.2015.12.010 (2016).

17. Gabay, C. & Kushner, I. Acute-Phase Proteins and Other Systemic Responses to Inflammation. New England Journal of Medicine 340, 448–454 (1999).

18. O’Hara, R., Murphy, E. P., Whitehead, A. S., FitzGerald, O. & Bresnihan, B. Acute-phase serum amyloid A production by rheumatoid arthritis synovial tissue. Arthritis Res 2, 142–144 (2000).

19. Abbas, A. A. et al. Role of Serum Amyloid A as a Biomarker for Predicting the Severity and Prognosis of COVID-19. J Immunol Res 2022, 6336556 (2022).

20. Meek, R. L., Eriksen, N. & Benditt, E. P. Murine serum amyloid A3 is a high density apolipoprotein and is secreted by macrophages. Proc Natl Acad Sci U S A 89, 7949–7952 (1992).

21. Shah, C., Hari-Dass, R. & Raynes, J. G. Serum amyloid A is an innate immune opsonin for Gramnegative bacteria. Blood 108, (2006).

22. Ye, R. D. & Sun, L. Emerging functions of serum amyloid A in inflammation. J Leukoc Biol 98, 923–929 (2015).

23. Frame, N. M. & Gursky, O. Structure of serum amyloid A suggests a mechanism for selective lipoprotein binding and functions: SAA as a hub in macromolecular interaction networks. FEBS Lett 590, (2016).

24. Bindu G H., Rao, V. S. & Kakkar, V. V. Friend turns foe: Transformation of anti-inflammatory hdl to proinflammatory HDL during acute-phase response. Cholesterol vol. 2011 Preprint at 10.1155/2011/274629 (2011).

25. Ogasawara, K. et al. A serum amyloid A and LDL complex as a new prognostic marker in stable coronary artery disease. Atherosclerosis 174, (2004).

26. den Hartigh, L. J., May, K. S., Zhang, X. S., Chait, A. & Blaser, M. J. Serum amyloid A and metabolic disease: evidence for a critical role in chronic inflammatory conditions. Frontiers in Cardiovascular Medicine vol. 10 1197432 Preprint at 10.3389/fcvm.2023.1197432 (2023).

27. Colón, W., Javier Aguilera, J. & Srinivasan, S. Intrinsic stability, oligomerization, and amyloidogenicity of HDL-free serum amyloid A. Adv Exp Med Biol 855, 117–134 (2015).

28. Lu, J., Yu, Y., Zhu, I., Cheng, Y. & Sun, P. D. Structural mechanism of serum amyloid A-mediated inflammatory amyloidosis. Proc Natl Acad Sci U S A 111, (2014).

29. Sorić Hosman, I., Kos, I. & Lamot, L. Serum Amyloid A in Inflammatory Rheumatic Diseases: A Compendious Review of a Renowned Biomarker. Frontiers in Immunology vol. 11 631299 Preprint at 10.3389/fimmu.2020.631299 (2021).

30. Chen, R. et al. Serum amyloid protein A in inflammatory bowel disease: from bench to bedside. Cell Death Discovery vol. 9 154 Preprint at 10.1038/s41420-023-01455-5 (2023).

31. Jahangiri, A. et al. Serum amyloid A is found on ApoB-containing lipoproteins in obese humans with diabetes. Obesity 21, (2013).

32. Zinellu, A., Paliogiannis, P., Carru, C. & Mangoni, A. A. Serum amyloid A concentrations, COVID-19 severity and mortality: An updated systematic review and meta-analysis. International Journal of Infectious Diseases vol. 105 668–674 Preprint at 10.1016/j.ijid.2021.03.025 (2021).

33. Tasar, S. et al. Serum Amyloid A Levels and Severity of COVID-19 in Children. Indian Pediatr 60, 217–220 (2023).

34. Almusalami, E. M., Lockett, A., Ferro, A. & Posner, J. Serum amyloid A—A potential therapeutic target for hyper-inflammatory syndrome associated with COVID-19. Frontiers in Medicine vol. 10 1135695 Preprint at 10.3389/fmed.2023.1135695 (2023).

35. Siman-Tov, R. et al. Elevated Serum Amyloid A Levels Contribute to Increased Platelet Adhesion in COVID-19 Patients. Int J Mol Sci 23, 14243 (2022).

36. Mizurini, D. M., Hottz, E. D., Bozza, P. T. & Monteiro, R. Q. Fundamentals in Covid-19-Associated Thrombosis: Molecular and Cellular Aspects. Frontiers in Cardiovascular Medicine vol. 8 785738 Preprint at 10.3389/fcvm.2021.785738 (2021).

37. Li, Y., Xiaojing, H., Zhuanyun, L., Li, D. & Yang, J. Prognostic value of serum amyloid A in COVID-19: A meta-analysis. Medicine (United States) vol. 101 e28880 Preprint at 10.1097/MD.0000000000028880 (2022).

38. Hammarström, P. & Nyström, S. Viruses and amyloids - a vicious liaison. Prion vol. 17 82–104 Preprint at 10.1080/19336896.2023.2194212 (2023).

39. Galkin, A. P. Hypothesis: AA amyloidosis is a factor causing systemic complications after coronavirus disease. Prion 15, 53–55 (2021).

40. Lockett, A. D., Wu, Y. & Gunst, S. J. Elastase alters contractility and promotes an inflammatory synthetic phenotype in airway smooth muscle tissues. Am J Physiol Lung Cell Mol Physiol 314, L626–L634 (2018).

41. Döring, G. The role of neutrophil elastase in chronic inflammation. In American Journal of Respiratory and Critical Care Medicine vol. 150 S114–S117 (1994).

42. Karampoor, S. et al. A possible pathogenic correlation between neutrophil elastase (NE) enzyme and inflammation in the pathogenesis of coronavirus disease 2019 (COVID-19). Int Immunopharmacol 100, 108137 (2021).

43. Charnley, M. et al. Neurotoxic amyloidogenic peptides in the proteome of SARS-COV2: potential implications for neurological symptoms in COVID-19. Nat Commun 13, 3387 (2022).

44. Nyström, S. & Hammarström, P. Amyloidogenesis of SARS-CoV-2 Spike Protein. J Am Chem Soc 144, 8945–8950 (2022).

45. Seth, P. & Sarkar, N. A comprehensive mini-review on amyloidogenesis of different SARS-CoV-2 proteins and its effect on amyloid formation in various host proteins. 3 Biotech vol. 12 322 Preprint at 10.1007/s13205-022-03390-1 (2022).

46. Semerdzhiev, S. A., Fakhree, M. A. A., Segers-Nolten, I., Blum, C. & Claessens, M. M. A. E. Interactions between SARS-CoV-2 N-Protein and α-Synuclein Accelerate Amyloid Formation. ACS Chem Neurosci 13, 143–150 (2022).

47. Sinha, N. & Thakur, A. K. Likelihood of amyloid formation in COVID-19-induced ARDS. Trends in Microbiology vol. 29 967–969 Preprint at 10.1016/j.tim.2021.03.008 (2021).

48. Jana, A. K., Greenwood, A. B. & Hansmann, U. H. E. Presence of a SARS-CoV-2 Protein Enhances Amyloid Formation of Serum Amyloid A. Journal of Physical Chemistry B 125, 9155–9167 (2021).

49. Nady, A., Reichheld, S. E. & Sharpe, S. Structural studies of a serum amyloid A octamer that is primed to scaffold lipid nanodiscs. Protein Science 33, e4982 (2024)

50. Beerten, J. et al. WALTZ-DB: A benchmark database of amyloidogenic hexapeptides. Bioinformatics 31, 1698–1700 (2015).

51. Conchillo-Solé, O. et al. AGGRESCAN: A server for the prediction and evaluation of ‘hot spots’ of aggregation in polypeptides. BMC Bioinformatics 8, (2007).

52. Gasteiger, E. et al. ExPASy: The proteomics server for in-depth protein knowledge and analysis. Nucleic Acids Res 31, 3784–3788 (2003).

53. Scheres, S. H. W. RELION: Implementation of a Bayesian approach to cryo-EM structure determination. J Struct Biol 180, 519–530 (2012).

54. Delaglio, F. et al. NMRPipe: A multidimensional spectral processing system based on UNIX pipes. J Biomol NMR 6, 277–293 (1995).

55. Surya, W., Li, Y. & Torres, J. Structural model of the SARS coronavirus E channel in LMPG micelles. Biochim Biophys Acta Biomembr 1860, 1309–1317 (2018).

56. Zhou, S. et al. SARS-CoV-2 E protein: Pathogenesis and potential therapeutic development. Biomedicine and Pharmacotherapy vol. 159 114242 Preprint at 10.1016/j.biopha.2023.114242 (2023).

57. Mustafa, Z., Zhanapiya, A., Kalbacher, H. & Burster, T. Neutrophil Elastase and Proteinase 3 Cleavage Sites Are Adjacent to the Polybasic Sequence within the Proteolytic Sensitive Activation Loop of the SARS-CoV-2 Spike Protein. ACS Omega 6, 7181–7185 (2021).

58. Schoeman, D. & Fielding, B. C. Coronavirus envelope protein: Current knowledge. Virology Journal vol. 16 69 Preprint at 10.1186/s12985-019-1182-0 (2019).

59. Kayed, R. et al. Annular protofibrils area structurally and functionally distinct type of amyloid oligomer. Journal of Biological Chemistry 284, 4230–4237 (2009).

60. Lasagna-Reeves, C. A. et al. The formation of tau pore-like structures is prevalent and cell specific: Possible implications for the disease phenotypes. Acta Neuropathol Commun 2, (2014).

61. Lasagna-Reeves, C. A., Glabe, C. G. & Kayed, R. Amyloid-β annular protofibrils evade fibrillar fate in Alzheimer disease brain. Journal of Biological Chemistry 286, 22122–22130 (2011).

62. Lattanzi, V. et al. Amyloid β 42 fibril structure based on small-angle scattering. Proc Natl Acad Sci U S A 118, e21122783118 (2021).

63. Infield, D. T. et al. Cation-π Interactions and their Functional Roles in Membrane Proteins: Cation-π interactions in membrane proteins. Journal of Molecular Biology vol. 433 167035 Preprint at 10.1016/j.jmb.2021.167035 (2021).

64. Mamsa, S. S. A. & Meloni, B. P. Arginine and Arginine-Rich Peptides as Modulators of Protein Aggregation and Cytotoxicity Associated With Alzheimer’s Disease. Frontiers in Molecular Neuroscience vol. 14 759729 Preprint at 10.3389/fnmol.2021.759729 (2021).

65. Das, M. & Gursky, O. Amyloid-forming properties of human apolipoproteins: Sequence analyses and structural insights. Adv Exp Med Biol 855, 175–211 (2015).

66. Janardhan, V., Janardhan, V. & Kalousek, V. COVID-19 as a Blood Clotting Disorder Masquerading as a Respiratory Illness: A Cerebrovascular Perspective and Therapeutic Implications for Stroke Thrombectomy. Journal of Neuroimaging vol. 30 555–561 Preprint at 10.1111/jon.12770 (2020).

67. Mir, T. H. et al. Post COVID-19 AA amyloidosis of the kidneys with rapidly progressive renal failure. Prion 17, (2023).

68. Fotuhi, M., Mian, A., Meysami, S. & Raji, C. A. Neurobiology of COVID-19. Journal of Alzheimer’s Disease vol. 76 Preprint at 10.3233/JAD-200581 (2020).

69. Onishi, T. et al. The balance of comprehensive coagulation and fibrinolytic potential is disrupted in patients with moderate to severe COVID-19. Int J Hematol 115, (2022).

70. Gazit, E. A possible role for π-stacking in the self-assembly of amyloid fibrils. The FASEB Journal 16, 77–83 (2002).

71. Stanković, I. M., Niu, S., Hall, M. B. & Zarić, S. D. Role of aromatic amino acids in amyloid self-assembly. International Journal of Biological Macromolecules vol. 156 949–959 Preprint at 10.1016/j.ijbiomac.2020.03.064 (2020).

72. Wang, Y. et al. Serum amyloid A 2.2 refolds into a octameric oligomer that slowly converts to a more stable hexamer. Biochem Biophys Res Commun 407, (2011).

73. Takase, H., Tanaka, M., Miyagawa, S., Yamada, T. & Mukai, T. Effect of amino acid variations in the central region of human serum amyloid A on the amyloidogenic properties. Biochem Biophys Res Commun 444, 492–497 (2014).

74. Patel, H., Bramall, J., Waters, H., De Beer, M. C. & Woo, P. Expression of recombinant human serum amyloid A in mammalian cells and demonstration of the region necessary for high-density lipoprotein binding and amyloid fibril formation by site-directed mutagenesis. Biochemical Journal 318, 1041–1049 (1996).

75. Jayaraman, S., Fändrich, M. & Gursky, O. Synergy between serum amyloid a and secretory phospholipase A2. Elife 8, e46630 (2019).

