## Supplementary for "An amyloidogenic fragment of the SARS CoV-2 envelope protein promotes serum amyloid A misfolding and fibrillization"

### Supplementary Information

| Protein | Sequence |
|---|---|
| **Spike** | MFVFLVLLPLVSSQCVNLTTRTQLPPAYTNSFTRGVYYPDKVFRSSVLHSTQDLFLPFFSNVTWFHAIHVSGTNGTKRFDNPVLPFNDGVYFASTEKSNIIRGWIFGTTLDSKTQSLLIVNNATNVVIKVCEFQFCNDPFLGVYYHKNNKSWMESEFRVYSSANNCTFEYVSQPFLMDLEGKQGNFKNLREFVFKNIDGYFKIYSKHTPINLVRDLPQGFSALEPLVDLPIGINITRFQTLLALHRSYLTPGDSSSGWTAGAAAYYVGYLQPRTFLLKYNENGTITDAVDCALDPLSETKCTLKSFTVEKGIYQTSNFRVQPTESIVRFPNITNLCPFGEVFNATRFASVYAWNRKRISNCVADYSVLYNSASFSTFKCYGVSPTKLNDLCFTNVYADSFVIRGDEVRQIAPGQTGKIADYNYKLPDDFTGCVIAWNSNNLDSKVGGNYNYLYRLFRKSNLKPFERDISTEIYQAGSTPCNGVEGFNCYFPLQSYGFQPTNGVGYQPYRVVVLSFELLHAPATVCGPKKSTNLVKNKCVNFNFNGLTGTGVLTESNKKFLPFQQFGRDIADTTDAVRDPQTLEILDITPCSFGGVSVITPGTNTSNQVAVLYQDVNCTEVPVAIHADQLTPTWRVYSTGSNVFQTRAGCLIGAEHVNNSYECDIPIGAGICASYQTQTNSPRRARSVASQSIIAYTMSLGAENSVAYSNNSIAIPTNFTISVTTEILPVSMTKTSVDCTMYICGDSTECSNLLLQYGSFCTQLNRALTGIAVEQDKNTQEVFAQVKQIYKTPPIKDFGGFNFSQILPDPSKPSKRSFIEDLLFNKVTLADAGFIKQYGDCLGDIAARDLICAQKFNGLTVLPPLLTDEMIAQYTSALLAGTITSGWTFGAGAALQIPFAMQMAYRFNGIGVTQNVLYENQKLIANQFNSAIGKIQDSLSSTASALGKLQDVVNQNAQALNTLVKQLSSNFGAISSVLNDILSRLDKVEAEVQIDRLITGRLQSLQTYVTQQLIRAAEIRASANLAATKMSECVLGQSKRVDFCGKGYHLMSFPQSAPHGVVFLHVTYVPAQEKNFTTAPAICHDGKAHFPREGVFVSNGTHWFVTQRNFYEPQIITTDNTFVSGNCDVVIGIVNNTVYDPLQPELDSFKEELDKYFKNHTSPDVDLGDISGINASVVNIQKEIDRLNEVAKNLNESLIDLQELGKYEQYIKWPWYIWLGFIAGLIAIVMVTIMLCCMTSCCSCLKGCCSCGSCCKFDEDDSEPVLKGVKLHYT |
| **Envelope** | MYSFVSEETGTLIVNSVLLFLAFVVFLLVTLAILTALRLCAYCCNIVNVSLVKPSFYVYSRVKNLNSSRVPDLLV |
| **Nucleocapsid** | MSDNGPQNQRNAPRITFGGPSDSTGSNQNGERSGARSKQRRPQGLPNNTASWFTALTQHGKEDLKFPRGQGVPINTNSSPDDQIGYYRRATRRIRGGDGKMKDLSPRWYFYYLGTGPEAGLPYGANKDGIIWVATEGALNTPKDHIGTRNPANNAAIVLQLPQGTTLPKGFYAEGSRGGSQASSRSSSRSRNSSRNSTPGSSRGTSPARMAGNGGDAALALLLLDRLNQLESKMSGKGQQQQGQTVTKKSAAEASKKPRQKRTATKAYNVTQAFGRRGPEQTQGNFGDQELIRQGTDYKHWPQIAQFAPSASAFFGMSRIGMEVTPSGTWLTYTGAIKLDDKDPNFKDQVILLNKHIDAYKTFPPTEPKKDKKKKADETQALPQRQKKQQTVTLLPAADLDDFSKQLQQSMSSADSTQA |
| **Matrix** | MADNGTITVEELKQLLEQWNLVIGFLFLAWIMLLQFAYSNRNRFLYIIKLVFLWLLWPVTLACFVLAAVYRINWVTGGIAIAMACIVGLMWLSYFVASFRLFARTRSMWSFNPETNILLNVPLRGTIVTRPLMESELVIGAVIIRGHLRMAGHSLGRCDIKDLPKEITVATSRTLSYYKLGASQRVGTDSGFAAYNRYRIGNYKLNTDHAGSNDNIALLVQ |

**Figure S1. The proteome of SARS-CoV-2 is concentrated with amyloidogenic sequences.** WALTZ (https:waltz.switchlab.org/) [1] amyloid prediction algorithm was used to predict segments of SARS-CoV-2 S, E, N, and M Proteins (Protein IDs: P0DTC2, P0DTC4, P0DTC9, P0DTC5) with high propensity (red) to form amyloid fibrils.

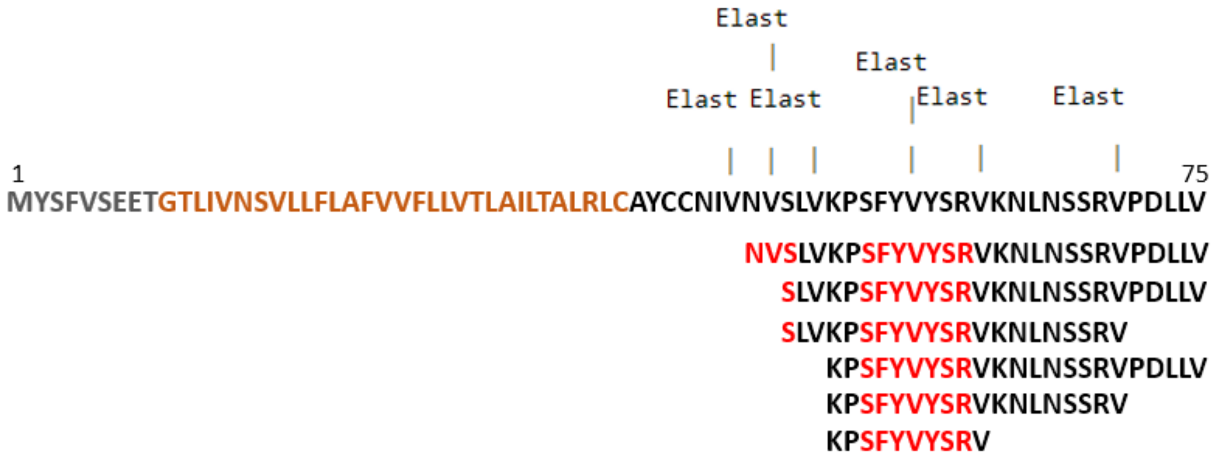


**Figure S2. Neutrophil elastase cleavage sites in the C-terminal domain (CTD) of the SARS-CoV-2 envelope protein. The sequence of the SARS-CoV-2 E protein is shown along with predicted** elastase cleavage sites (Expasy PeptideCutter, https://web.expasy.org/peptide_cutter/) [2]. The segments predicted to be amyloidogenic by WALTZ prediction algorithm are highlighted in red in the possible peptide products. The CTD sequence is shown in black, the transmembrane domain in gold, and N-terminal domain in grey.

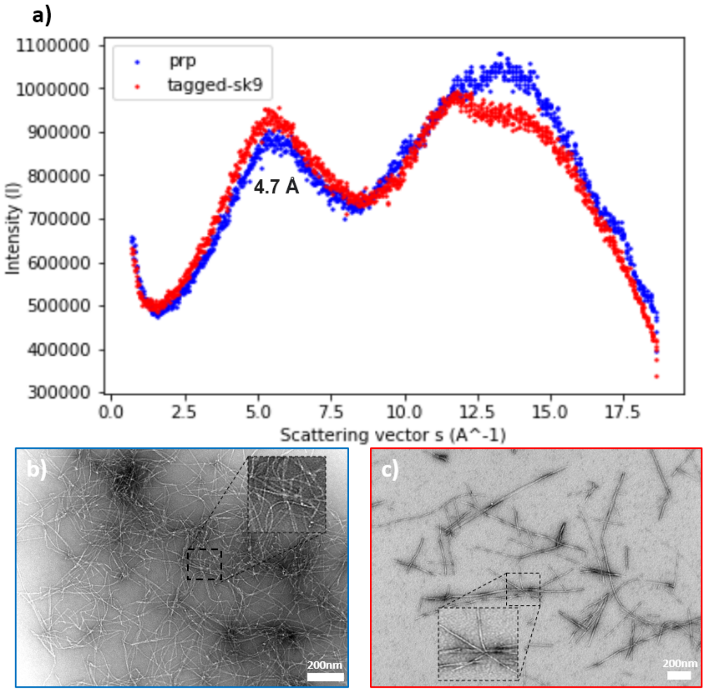


**Figure S3. Wide-angle X-ray scattering (WAXS) and transmission electron microscopy (TEM) of tagged-SK9 and PrP(244-249) fibrils.** (a) WAXS scattering of fibrils formed by an amyloidogenic fragment of the human prion protein, PrP(244-249) [3] (blue) and fibrils of dansyl-SK9 (red) at 25 °C. Both display an identical Bragg peak representative of the interstrand hydrogen bond distance of 4.7 Å, and similar peaks representative of their inter-sheet packing distance (11-14 Å). TEM images of (b) PrP(244-249) fibrils (magnification = 40000 x) and (c) dansyl-SK9GS fibrils (magnification = 30000 x). TEM samples were stained with 2% uranyl acetate.

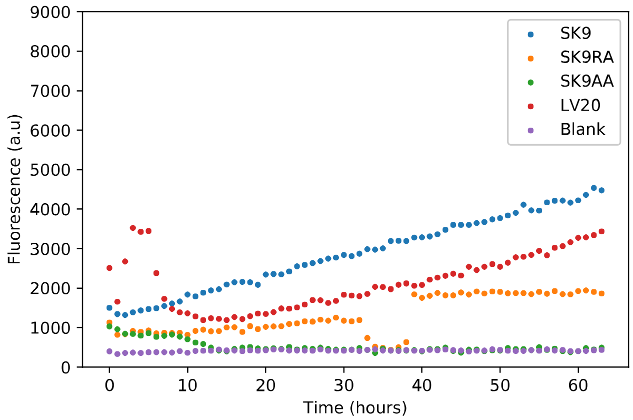


**Figure S4. Thioflavin T fluorescence assay on SK9 and related peptides.** All peptides were dissolved in PBS buffer at a final concentration of 0.05 mg/mL. ThT was added at time 0 and fluorescence emission at 485 nm (excitation at 450 nm) were recorded as a function of incubation time. All experiments were performed at 37 °C.

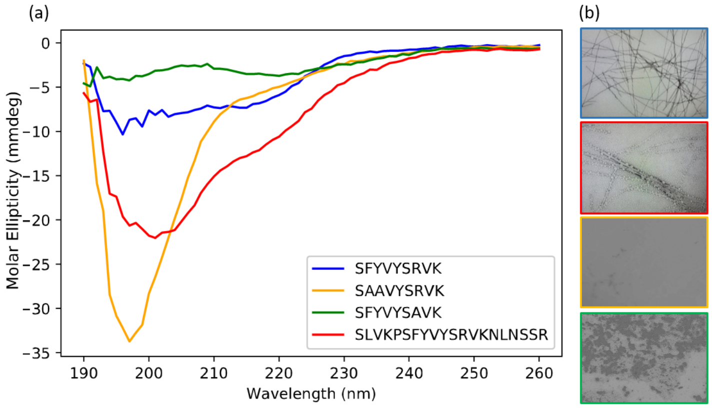


**Figure S5. Circular dichroism (CD) spectra and TEM images of the peptides derived from the SARS-CoV-2 envelope protein CTD**. (a) CD spectra are shown for peptides SK9 (blue), SK9AA (yellow), SK9RA (green), and LV20 (red), with sequences as shown. All CD experiments were performed at room temperature and peptides were immediately dissolved in 20 mM Tris (pH 8) prior to measurements. (b) Negative-stain TEM images (magnification = 40000x) are shown for the corresponding peptides (SK9 (blue), SK9AA (yellow), SK9RA (green), and LV20 (red)) following 3 days of incubation at 37 °C.

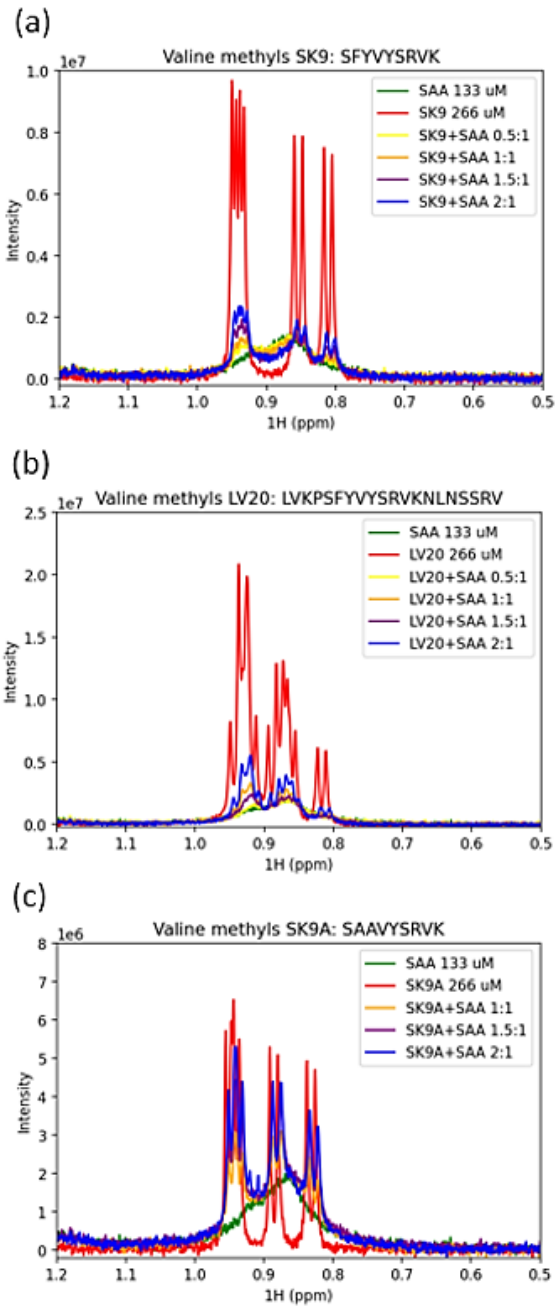


**Figure S6. NMR data confirm that SK9 and LV20 interact more strongly with SAA than SK9A.** Solution $^1$H NMR spectra (showing only the valine methyl region) are presented for (a) SK9, (b) LV20 and (c) SK9AA in the presence and absence of different ratios of SAA. Control spectra of SAA only are also shown in each panel. SK9RA was not used in the experiments.

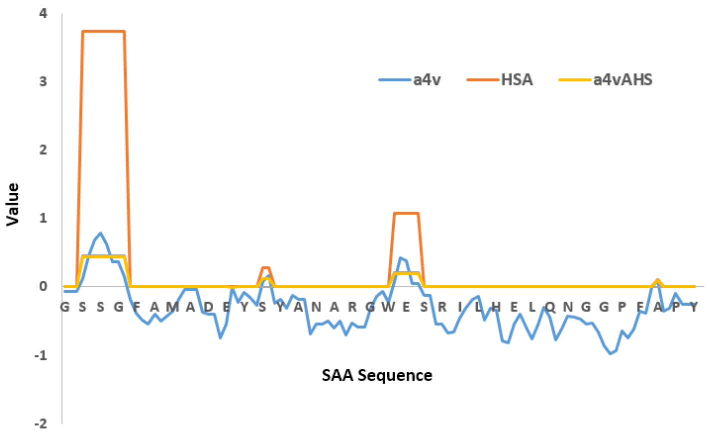


**Figure S7. AGGRESCAN (http:bioinf.uab.es/aggrescan/) [4] results predicting aggregation hotspots in the sequence of SAA.** The window average of aggregation propensity (a4v, blue) is calculated from individual amino acid propensities. The algorithm identifies hotspots as consecutive amino acids with a4v values falling above a given threshold, with average values within a hotspot plotted as a4vAHS (yellow). Hotspots are scored using a hotspot area value (HAS, red).

### References


1. Beerten, J. *et al.* WALTZ-DB: A benchmark database of amyloidogenic hexapeptides. *Bioinformatics* **31**, 1698–1700 (2015).
2. Gasteiger, E. *et al.* Protein identification and analysis tools in the ExPASy server. *Methods Mol Biol* **112**, 531–552 (1999).
3. Yau, J. & Sharpe, S. Structures of amyloid fibrils formed by the prion protein derived peptides PrP(244-249) and PrP(245-250). *J Struct Biol* **180**, 290–302 (2012).
4. Conchillo-Solé, O. *et al.* AGGRESCAN: A server for the prediction and evaluation of 'hot spots' of aggregation in polypeptides. *BMC Bioinformatics* **8**, (2007).
